# Viscoelasticity and cell jamming state transition

**DOI:** 10.1101/2021.03.19.436195

**Authors:** Ivana Pajic-Lijakovic, Milan Milivojevic

## Abstract

Although collective cell migration (CCM) is a highly coordinated migratory mode, perturbations in the form of jamming state transitions and vice versa often occur even in 2D. These perturbations are involved in various biological processes, such as embryogenesis, wound healing and cancer invasion. CCM induces accumulation of cell residual stress which has a feedback impact to cell packing density. Density-mediated change of cell mobility influences the state of viscoelasticity of multicellular systems and on that base the jamming state transition. Although a good comprehension of how cells collectively migrate by following molecular rules has been generated, the impact of cellular rearrangements on cell viscoelasticity remains less understood. Thus, considering the density driven evolution of viscoelasticity caused by reduction of cell mobility could result in a powerful tool in order to address the contribution of cell jamming state transition in CCM and help to understand this important but still controversial topic. In addition, five viscoelastic states gained within three regimes: (1) convective regime, (2) conductive regime, and (3) damped-conductive regime was discussed based on the modeling consideration with special emphasis of jamming and unjamming states.

## 1. Introduction

Main features of cell rearrangement during CCM related to the viscoelasticity of multicellular systems are essential for deeper understanding of various biomedical processes such as wound healing, morphogenesis, and tumorigenesis (Blanchard et al., 2019; Barriga et al., 2018; Barriga and Mayor, 2019; Petrungaro et al., 2019; Pajic-Lijakovic and Milivojevic, 2019a,2020a). The viscoelasticity of multicellular systems depends on the configuration of migrating cells and the rate of its change occurred by migrating-to-resting cell state transitions and vice versa (Pajic-Lijakovic and Milivojevic, 2019b,c). These transitions have been considered as the cell jamming state transitions (Angelini et al., 2011; Nnetu et al., 2013; Garcia et al., 2015; Bi et al., 2015; Park et al., 2016; Oswald et al. 2017). Several inter-dependent factors influence the jamming state transitions such as: (1) an increase in cell packing density (Henkes et al., 2011; Nnetu et al., 2013), (2) cell−cell adhesion energy (Garcia et al., 2015; Bi et al., 2015), (3) magnitude of cellular forces and persistence time for these forces (Garcia et al., 2015), (3) cell shape (Garcia et al., 2015; Park et al., 2016), (3) contact inhibition of locomotion (CIL) (Zimmermann et al., 2016). The reduction in cell mobility influences the state of viscoelasticity and on that base the cell jamming state transition. Despite extensive research devoted to study of the cell jamming, we still do not understand the process from the standpoint of rheology. However, the mechanism or mechanisms through which this jamming state comes about are connected with the viscoelasticity (Pajic-Lijakovic and Milivojevic, 2019c).

The change of the state of viscoelasticity caused by reducing the movement of system constituents has been considered for various soft matter systems such as: (1) granular systems and micro-gels (Edwards and Grinev, 1998; Edwards, 2005; Ciamarra et al., 2006), (2) entangled polymer systems and coacervate based systems (Liu et al., 2006; Liu et al., 2017); (3) polymer nano-composites (Hassanabadi et al., 2014). The movement reduction can be induced: (1) by increasing frequency in the vibrational flow field or decreasing the observation time, (2) by increasing the concentration of the system constituents and their rigidity, and (3) by decreasing the temperature (i.e. the glass state transition) (Pajic-Lijakovic, 2021). The apparent increase in the rigidity of the system constituents can be induced by intensifying the field of interactions quantified by the concept of excluded volume (Neal et al., 1998). Liu et al. (2017) considered the rheological response of coacervate-based systems in oscillatory shear field by increasing frequency and distinguished four regimes: (1) low-frequency terminal regime described by the Maxwell model (a viscoelastic liquid), (2) middle-frequency plateau regime described by the Kelvin-Voigt model (a viscoelastic solid), (3) higher-frequency transition regime (a viscoelastic solid), and (4) high-frequency jamming regime (a viscoelastic solid). The terminal regime is characterized by intensive and disordering motion of the system constituents and intra-chain interactions which induces significant energy dissipation. Plateau regime corresponds to intensive entropic effects caused by inter-chain interactions (Pajic-Lijakovic, 2021). Transition regime corresponds to reduced mobility of the system constituents and accounts for local conformation changes of the chain parts described by entanglement relaxation time (Liu et al., 2006). In the case of polymer nanocomposites, the addition of nanoparticles to polymer systems intensifies their inter-connectivity and on that base reduces the mobility of polymer chains. The critical concentration of nanoparticles induces the jamming state transition (Hassanabadi et al., 2014). The main characteristics of the jamming state are: (1) migration of the system constituents is much damped such that the diffusion coefficient tends to zero, (2) relaxation time tends to infinity, (3) storage modulus *G* ^’^(*ω*) and loss modulus *G*^”^ (*ω*) satisfy the condition *G* ^’^ (*ω*)/ *G* ^’’^ (*ω*) = *const* > 1 (Honter and Weeks, 2012; Pajic-Lijakovic and Milivojevic, 2019c).

The density-driven cell jamming state transition during CCM has been widely studied (Angelini et al., 2011; Nnetu et al., 2013; Garcia et al., 2015; Bi et al., 2015; Park et al., 2016; Oswald et al. 2017). However, the corresponding change of the state of viscoelasticity has not taken into consideration. To fill this gap, we provide a systematic theoretical consideration from the standpoint of rheology. Angelini et al. (2011) pointed out to the density-driven transition from convective to the conductive mechanism of CCM. This transition influences the state of viscoelasticity. Garcia et al. (2015) considered the velocity correlation length as function of cell velocity. They revealed that the correlation length (1) increases with cell velocity for the cell velocities lower than 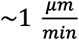 and (2) decreases with cell velocity for the cell velocities higher than 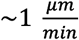. These two trends of cell movement correspond to various states of viscoelasticity. Further increase in the cell packing density leads to the reduction in cell mobility, manifested as a sub-diffusional cell migration, which induces the anomalous nature of energy dissipation (Nnetu et al., 2013). Consequently, density-driven cell rearrangement should be considered within three regimes: (1) convective regime, (2) conductive regime, and (3) damped-conductive regime which accounts for the cell jamming state. Every regime corresponds of several states of viscoelasticity. Garcia et al. (2015) pointed out that cell monolayers behave as amorphous solids under reduced cell velocities. Every viscoelastic state per regime should be characterized by appropriate stress-strain constitutive model.

Pajic-Lijakovic and Milivojevic (2019a,2020a) reported that CCM induces generation of stress, its relaxation, and the residual stress accumulation. The stress can be normal (compressive and tensile) and shear. Normal stress is primarily accumulated within a core region of migrating cell clusters during their movement through the dense environment (Pajic-Lijakovic and Milivojevic, 2019a;2021) and during collisions of migrating cell fronts (Nnetu et al., 2012). The normal residual stress accumulation is responsible for the increase in cell packing density (Trepat et al., 2009) and reduction of cell mobility (Pajic-Lijakovic and Milivojevic, 2021) and on that base for the jamming state transition (Pajic-Lijakovic, 2021). The theoretical consideration accounts for: characterization of the established regimes by cell velocities, the rate of cell packing density change and the viscoelasticity at a supracellular level. An attempt here is made to clarify the density-driven evolution of the viscoelasticity and the cell jamming state transition and to point out to the parameters which control the dynamics of cell long-time rearrangement and their inter-relations. Consequently, five states of viscoelasticity will be considered within three regimes by formulating various constitutive models.

## 2. Viscoelastic regimes change by increase in cell packing density

CCM induces generation of stress, its relaxation, and the residual stress accumulation (Pajic-Lijakovic and Milivojevic, 2019a;2020a). Distribution of normal (compressive and tensile) and shear residual stresses has been measured for 2D multicellular systems under *in vitro* conditions (Serra-Picamal et al., 2012; Tambe et al., 2013; Notbohm et al., 2016). Tambe et al. (2013) reported that the maximum stress corresponds to *100-150 Pa*. The normal residual stress accumulation induces an increase in the cell packing density (Trepat et al., 2009). An increase in cell packing density induces a decrease in cell velocity (Angelini et al., 2011; Nnetu et al., 2013; Tlili et al., 2018) and on that base a change the state of viscoelasticity. The cause-consequence relation was presented in Figure 1.

**Figure 1.**
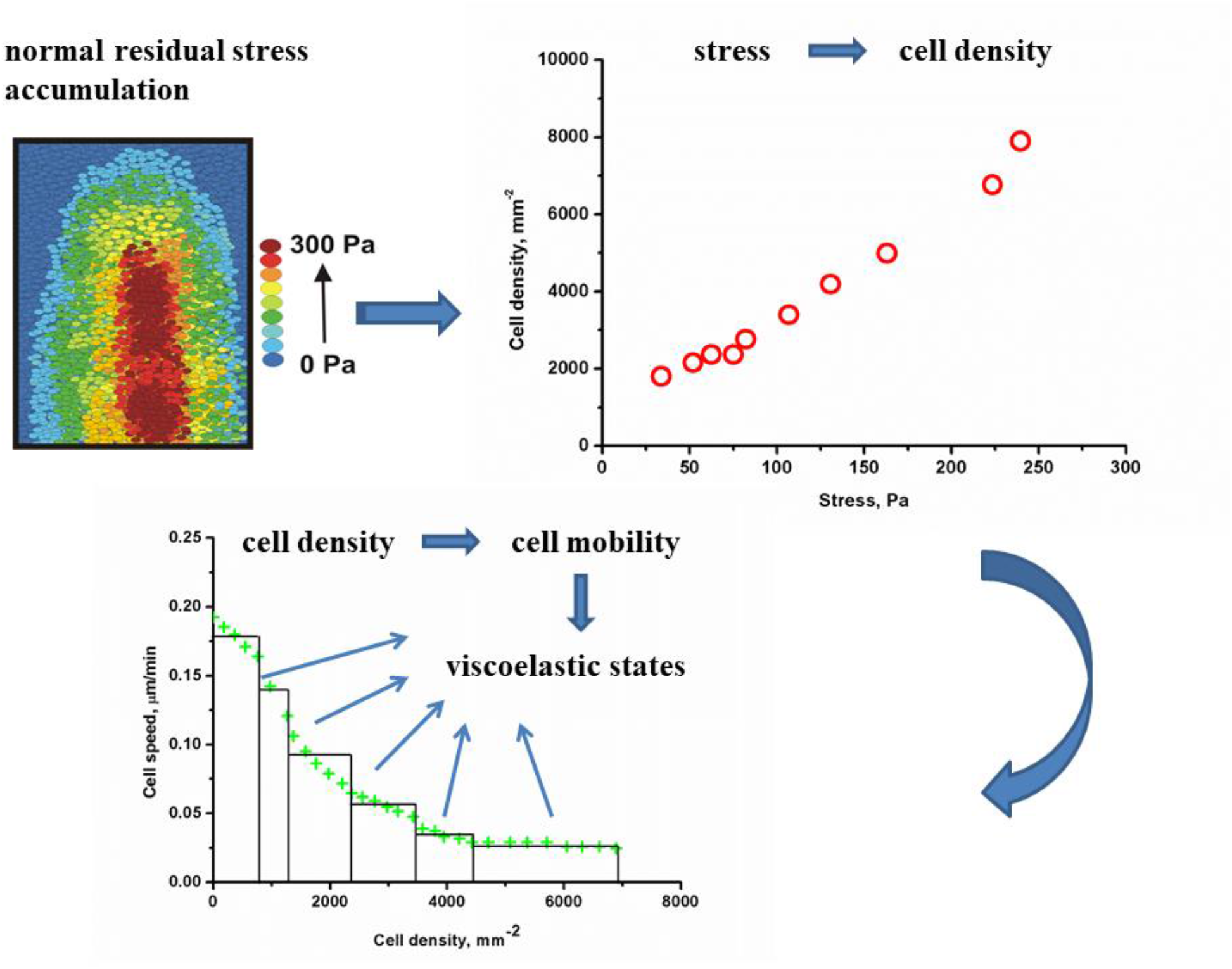
The cause-consequence relation between the normal residual stress accumulation, cell packing density, cell movement, and the states of viscoelasticity of 2D multicellular systems.

We considered here two scenarios of increase in cell packing density (1) within migrating clusters (labeled by A) caused by normal residual stress accumulation during their movement through dense environment and (2) caused by collision of velocity fronts (labeled by A and B) proposed by Pajic-Lijakovic and Milivojevic (2019a). The cell division is neglected at this time-scale. Both scenarios can lead to migrating-to-resting cell state transition (Pajic-Lijakovic and Milivojevic, 2019c) as was schematically presented in Figure 2.

**Figure 2.**
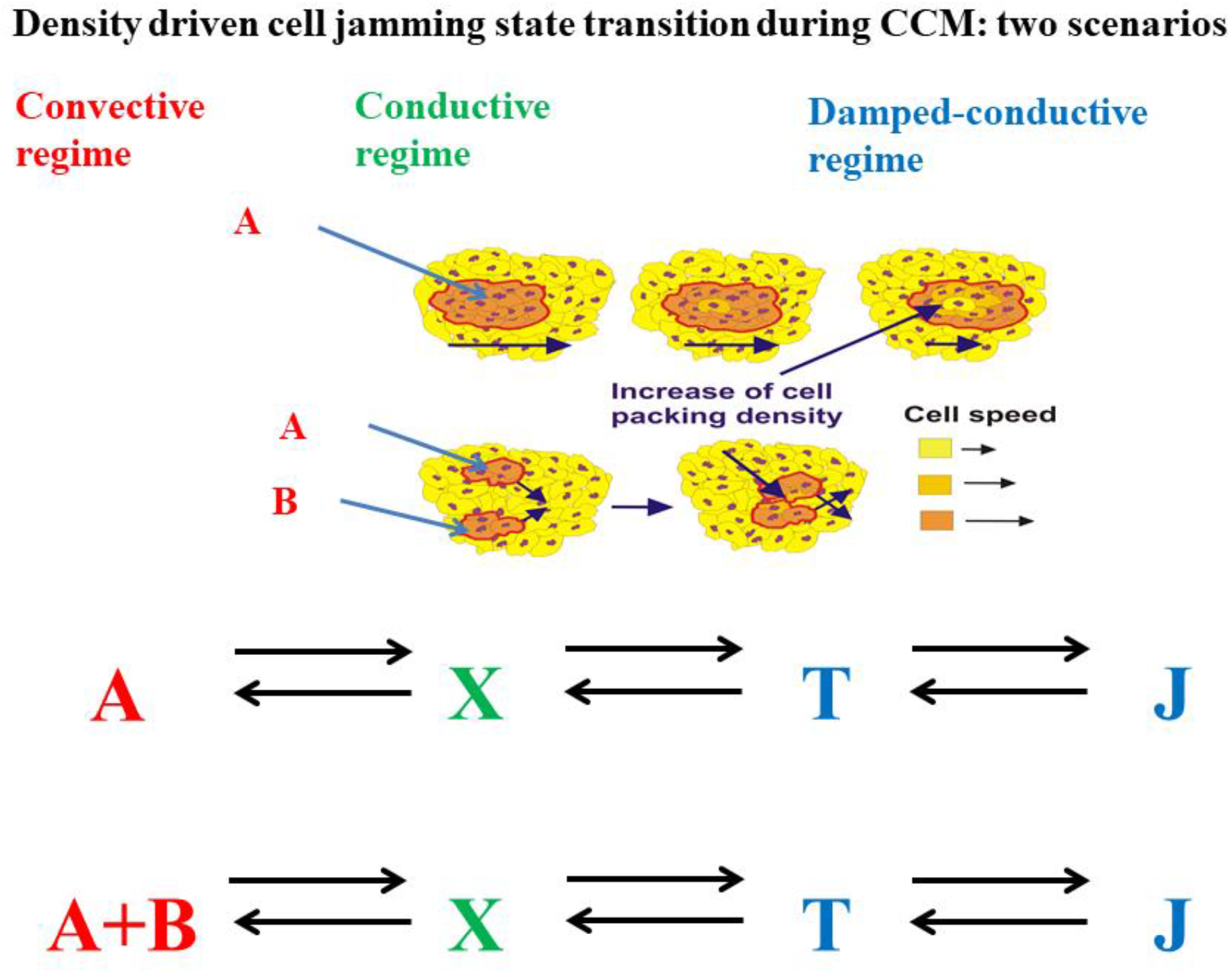
Two scenarios of the density-driven jamming state transition during CCM

The A and B represents various cell migrating states within a convective regime, X accounts for cell migration within the conductive regime, T represents the cell transient state occurred within the damped-conductive regime, and J represents the cell jamming state also occurred within the damped-conductive regime. Braumgarten and Tighe (2017) reported that systems have to pass through a transient state before reaching the jamming state. The underlying migration mode for the cell states labeled by A and B is convective, while the transient state X accounts for cell movement by conductive mechanism. Consequently, three regimes have been experimentally observed during CCM of cell monolayers such as: (1) convective cell regime, (2) conductive regime, and (3) damped-conductive regime which account for anomalous nature of energy dissipation (Angelini et al., 2011; Nnetu et al., 2013; Garcia et al., 2015). The transient and jamming states represent parts of this damped-conductive regime. Every regime will be characterized by cell velocity (i.e. the rate of change the cell displacement field caused by CCM), cell packing density, and the corresponding constitutive stress-strain relation. Transitions from the convective to the conductive regime and from conductive to the damped-conductive regime are induced by normal residual stress accumulation accompanied with an increase in cell packing density (Pajic-Lijakovic, 2021). On the other hand, feedback transitions from the damped-conductive to the conductive regime and from the conductive to the convective regime are caused by contact inhibition of locomotion (CIL) (Zimmermann et al., 2016; Alert and Trepat, 2020). CIL is responsible for the disordering of multicellular systems and weakening of cell-cell adhesion contacts (Mayor and Carmona-Fontaine, 2010). Density-driven transitions from a regime to regime could be treated as the viscoelastic phase transitions.

An increase in cell packing density leads to a decrease in effective volume per single cell and on that base intensifies cell-cell interactions. This effective volume can be expressed as:

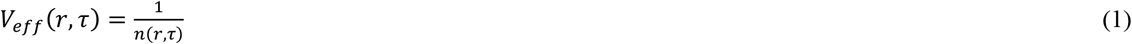

where *τ* is the long-time scale of several tens of minutes to hours, *n*(*r,τ*) is the local cell packing density equal to *n*(*r, τ*) *∑*_*i*_*δ*(*r* − *r*_*i*_), *V*_*eff*_(*r, τ*) is the effective volume per single cell equal to *V*_*eff*_ (*r, τ*) =*V*_*c*_ *+V*_*f*,_ *V*_*c*_, is the average single-cell volume, and *V*_*f*_ is the free volume per single cell. The increase in cell packing density induces a decrease in the free volume *V*_*f*_, while the single-cell volume stays approximately constant during CCM. The jamming state occurs when:

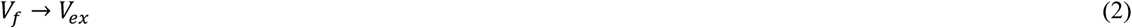

where *V*_*ex*_ is the excluded volume per single cell. This condition intensifies the CIL which can induce unjamming (Alert and Trepat, 2020). The concept of excluded volume has been introduced in physics in order to describe multi body interactions within various soft matter systems made by polymers, as well as, soft and rigid particles (Neal et al., 1998). The excluded volume depends on the interaction field and has been expressed from the second virial coefficient (Neal et al., 1998). We applied this concept to dense cell populations in order to define the minimum effective volume per single cell which corresponds to the jamming state and on that base the maximum cell packing density. The second virial coefficient is equal to:

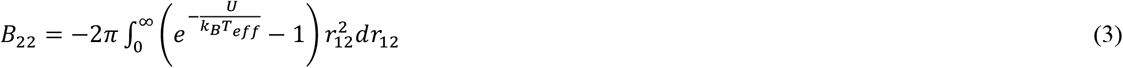

where *U* (*r*_12_) is the interaction potential between two cells, k_*B*_ is the Boltzmann constant, *T*_*eff*_ is the effective temperature. The cell potential U accounts for a short-range repulsion and mid-range attraction (Alert and Trepat, 2020). The repulsion has been modeled by hard-core or soft-core potentials (Smeets et al., 2016). The mid-range attraction accounts for cell-cell adhesion. When the potential *U*(*r*_12_) → ∞, the second virial coefficient tends to the cell excluded volume, i.e. *B*_22_ → *V*_*ex*_ (Neal et al., 1998). Consequently, the cell packing density for the jamming state *n*_*J*_ can be expressed as:

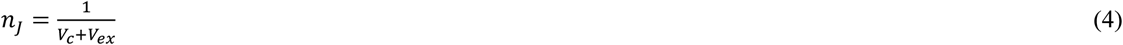

Where *V*_*eff min*_ = *V*_*c*_ + *V*_*ex*_ is the minimum of the effective volume per single cell. The value of *n*_*J*_ as well as *V*_*ex*_ varies with cell type and experimental conditions. Consequently, multicellular systems with similar densities can either remain fluid or jammed (Chen et al., 2018). For 2D CCM, the cell packing density at jamming is 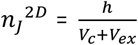 (where *h* is the average height per single cell). Consequently, the cell packing density increase (1) influences the cell migration state and on that base the single-cell shape (Nnetu et al., 2013), (2) influences the state of cell-cell and cell-matrix adhesion contacts (Garcia et al., 2015), and (3) intensifies CIL (Mayor and Carmona-Fontaine, 2010). In further consideration it is necessary to define a change of cell packing density with time for various established viscoelastic regimes.

The concept of effective temperature has been applied for considering rearrangement of various thermodynamic systems from glasses and sheared fluids to granular systems (Casas-Vazquez and Jou, 2003). Pajic-Lijakovic and Milivojevic (2019c;2021) applied this concept to a long-time rearrangement of dense cellular systems. The effective temperature, in this case, represents a product of cell mobility and has been expressed as 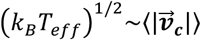 (where 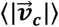 is the cell average speed) (Pajic-Lijakovic and Milivojevic, 2021). The effective temperature decreases from the convective to the damped-conductive regime. In further consideration, it is necessary to define the rate of packing density change per regime as a consequence of volumetric strain energy accumulation.

### 3. The cell packing density change for various regimes

A long-time change of cell packing density was expressed by modifying the model proposed by Murray et al. (1988):

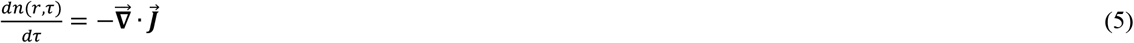

where *τ* is the long-time scale, 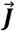 is the flux of cells equal to 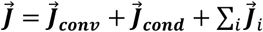, such that 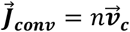 is the convective flux, 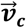 is cell velocity, 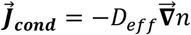 is the conductive flux, *D*_*eff*_ is the effective diffusion coefficient, while 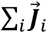 are the sum of various fluxes such as haptotaxis, galvanotaxis, chemotaxis, durotaxis and plithotaxis (Murray et al., 1988; Pajic-Lijakovic and Milivojevic, 2020c). All mentioned fluxes influences the rate 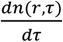.

The rate 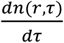 can be 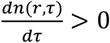 for convective and conductive regimes such that 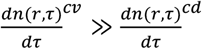 and 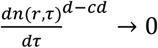 for the damped-conductive regime (where 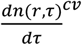 corresponds to the convective regime, 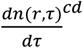 corresponds to the conductive regime, and 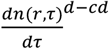 corresponds to the damped-conductive regime). The influence of: convective, conductive and damped-conductive modes of change the cell packing density on viscoelasticity of multicellular systems will be discussed in further modeling consideration.

The cell packing density change, for the convective regime (labeled by A and B in Figure 2), is expressed as:

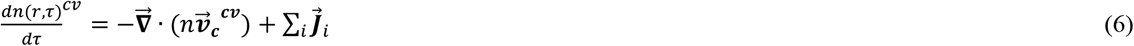

Where 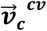 is the cell velocity for the convective regime. The cell packing density change, for the conductive regime (labeled by X) is:

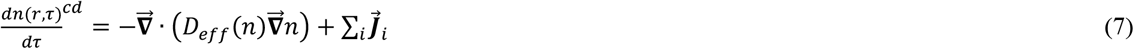

where *D*_*eff*_(*n*) is the effective diffusion coefficient. The effective diffusion coefficient decreases from 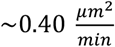 to 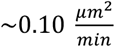 when the packing density of MDCK cells increases from 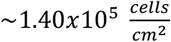 to 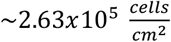 (Angelini et al., 2011). Rieu et al. (2000) reported that the diffusion coefficient for collectively migrated endodermal cells is 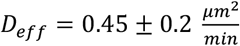, while for ectodermal cells the diffusion coefficient is 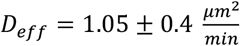. The cell velocity for the conductive regime can be expressed as:

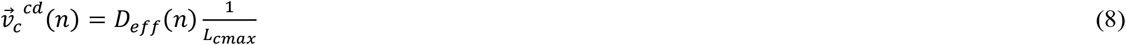

where *L*_*cmax*_ is the maximum velocity correlation length. Garcia et al. (2015) considered a change of the velocity correlation length with cell velocity. They distinguished two functional trends: (1) the *L*_*c*_ increases with the cell velocity and reaches the maximum value *L*_*cmax*_ for 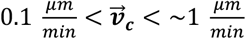 and (2) the *L*_*c*_ decreases with the cell velocity for for 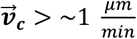. The correlation length slightly decreases with the cell packing density (Garcia et al., 2015). This decrease represents a consequence of a decrease in the diffusion coefficient (Angelini et al., 2011). The maximum correlation length for 2D CCM is *L*_*max*_ ∼ 150 *μm* (Petroli et al., 2021). Corresponding value of the velocity 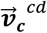 is in the range of 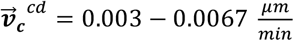 (eq. 8).

Consequently, two velocity trends distinguished by Garcia et al. (2015) correspond to the convective regime rather than the conductive regime. Serra-Picamal (2012) considered free expansion of Madin-Darbvy canine kidney type II cell (MDCK) monolayers on polyacrylamide gel and reported that the maximum velocity is 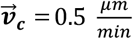. Notbohm et al. (2016) considered long-time rearrangement of confluent MDCK cell monolayers on polyacrylamide gel. They measured the maximum radial velocity during cell swirling motion equal to 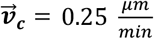. The cell velocities in both cases are significantly higher than the calculated conductive cell velocity 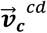. Clark and Vignjevic (2015) reported that the velocity of migrating cell clusters is 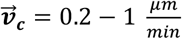 under *in vivo* conditions. Garcia et al. (2015) considered 2D CCM of various types of cells and pointed out that cell packing density increase from 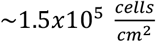 to 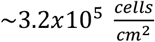 causes a decrease in the cell velocity for an order of magnitude. Tlili et al. (2018) considered density driven CCM of MDCK cell monolayers. They revealed that the cell packing density increase from 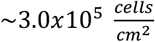 to 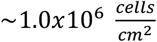 leads to a decrease in the cell velocity from 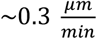 to zero. Nnetu et al. (2012) considered CCM of MCF-10A cell monolayers and obtained the maximum local cell velocity of 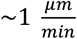 for the low cell packing density of 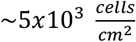. For the higher cell packing density of 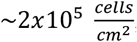, the maximum local cell velocity drops to 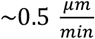 (Nnetu et al., 2012). Both cell velocities correspond to the convective regime. Further increase in cell packing density to 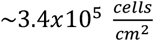, leads to a significant decrease in the cell velocity (Nnetu et al., 2013).

Nnetu et al. (2013) pointed out to the anomalous nature of energy dissipation during a cell rearrangement occurred under this condition (the damped-conductive regime). Cells undergo the sub-diffusion movement described by fractional derivatives (Podlubny, 1999). The corresponding minimum order of the fractional derivative *α* (i.e. the damping coefficient) is *α*∼0.2 (Nnetu et al., 2013). Within the damped-conductive regime cells passed from the transient state (labeled by T in Figure 2) for the damping coefficient equal to *α=* 1/2 (Baumgarten and Tighe, 2017) to the jamming state (labeled by J in Figure 2) for 0 < *α* < 1/2 during several hours (Nnetu et al., 2013; Pajic-Lijakovic, 2021). The prerequisite for unjamming state transition is the weakening of cell-cell adhesion contacts (Park et al., 2015). This weakening can be induced by CIL, which produce a repulsive force among the cells upon contact (Lin et al., 2018; Alert and Trepat, 2020). The cell packing density change, for the damped-conductive regime is:

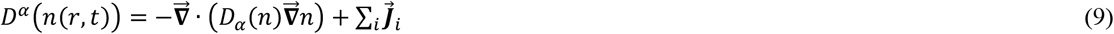

where *D*_*α*_ (*n*) is the damped-conductive diffusion coefficient which has the unit 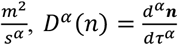 is the fractional derivative, and is the order of fractional derivative i.e. the damping coefficient of system structural changes which satisfies the condition *α* ≤ 1/2 (the transient and jamming states). Caputo’s definition of the fractional derivative of a function *n*(*r, τ*) was used, and it is given as (Podlubny, 1999):

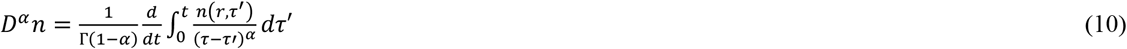

where Г(1− *α*) is a gamma function. The velocity for damped-conductive mechanism can be expressed as:

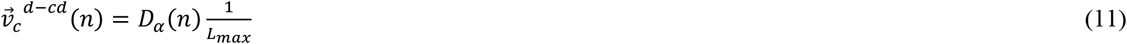

where *D*_*α*_(*n*) is the damped-conductive diffusion coefficient which has the unit 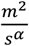, *α* is the damping coefficient such that *α* ≤*=* 1/2. The velocity for damped-conductive mechanism is 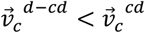. The main condition of the jamming state is that diffusion coefficient *D*_*α*_(*n*) → 0 (Hunter and Weeks, 2012).

The cell velocity at a mesoscopic level represents a rate of change the cell displacement field during CCM. For convective and conductive regimes it can be expressed as (Pajic-Lijakovic and Milivojevic, 2020c;2020b):

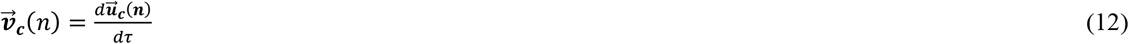

Where 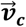 is convective velocity 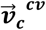 or conductive velocity 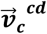 and 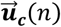 is the local cell displacement field equal to 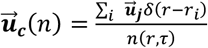 (where 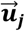 is the displacement field of the j-th cell). The cell velocity for the damped-conductive regime can be expressed as:

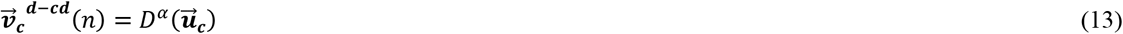

Where 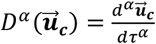 is the fractional derivative described by eq. 10. Volumetric and shear strains and the rate of their change can be expressed from the cell displacement field (Pajic-Lijakovic and Milivojevic, 2020b). Several factors influence the viscoelasticity of multicellular systems within various regimes such as (1) the rate of strain changes, (2) ability of stress to relax under constant strain condition, and (3) ability of stress to relax under constant strain rate condition (Pajic-Lijakovic and Milivojevic, 2017; 2019c; Pajic-Lijakovic, 2021). Further rheological consideration is accompanied with discussion of the free energy of cell rearrangement change as a function of: (1) viscoelasticity expressed by the volumetric strain energy density change, (2) effective temperature, (3) internal entropy production, (4) effective chemical potential, and (5) the rate of change the cell packing density.

## 4. Free energy of cell rearrangement

Free energy of cell rearrangement *F*_*rear*_ changes by changing the state of viscoelasticity. The rate of *F*_*rear*_ change: (1) for convective and conductive regimes is equal to 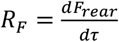 and (2) for the damped-conductive regime is equal to *R*_*F*_ = *D*^*α*^*F*_*rear*_ (where *D*^*α*^ (·) is the fractional derivative). The rate *R*_*F*_ depends on the change the strain energy density *R*_*W*,_ the effective tempereture, the internal entropy generation *R*_*S*_, the effective chemical potential 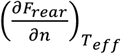, and the rate of change the cell packing density *R*_*n*_. The rate *R*_*F*_ is expressed as:

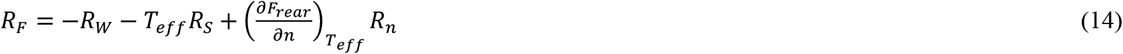

where *R*_*W*_ is the rate of change the volumetric strain energy density and *R*_*S*_ is the rate of increase the internal entropy. The rate *R*_*W*_ is equal to 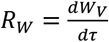 for convective and conductive regimes, while for the damped-conductive regime is equal to *R*_*W*_ = *D*^*α*^*W*_*V*_ (where 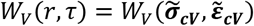 is the volumetric strain energy density, 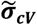 is the normal residual stress, 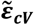 is the volumetric strain, and *D*^*α*^(·) is the fractional derivative). Consequently, the rate *R*_*W*_ depends on the state of viscoelasticity of multicellular system. The rate *R*_*n*_ is expressed by eq. 6 for the convective regime, eq. 7 for the conductive regime, and eq. 9 for the damped-conductive regime. The rate *R*_*S*_ is equal to 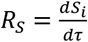 for convective and conductive regimes, while for the damped-conductive regime is equal to *R*_*S*_ *D*^*α*^*S*_*i*_ (where *S*_*i*_(*r, τ*) is the internal entropy). The internal entropy has expressed as (Pajic-Lijakovic and Milivojevic, 2020b;2021):

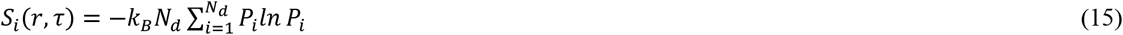

where *N*_*d*_ is the number of cells within the regime and *P*_*i*_ is the Boltzmann probability equal to 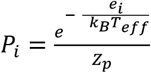, *Z*_*p*_ is the canonical partition function, *e*_*i*_ is the mechanical energy of single-cell. The mechanical energy *e*_*i*_ consists of active and passive parts (Koride et al., 2018). The passive part *e*_*pi*_ is *e*_*pi*_ = *γ*_*i*_(*A*_*i*_ − *A*_*0*_) *+ ∑*_*j*_ *Λl*_*ij*_, *γ*_*i*_ is the single-cell contribution to the tissue surface tension equal to 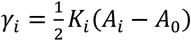, *K*_*i*_ is an effective bulk modulus of the cell, *A*_0_ is the reference area of cell while, *A*_*i*_ is the current area of the i-th cell such that *A*_*i*_ = *A*_*i*_(*ρ*_*A*_), *Λ* is the adhesion energy per unit length, *l*_*ij*_ is the edge length between vertex *i* and *j, U*_*a*_ is the active part of mechanical energy equal to 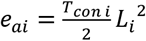 (where *T*_*con i*_ is the contractility coefficient, *L*_*i*_ and is the perimeter of the i-th cell) (Koride et al., 2018).

Every state of viscoelasticity corresponds to the current equilibrium state expressed by *R*_*F*_ = 0 and the free energy of rearrangement is *F*_*rear*_ ≈ *const*. Every current equilibrium state satisfies following conditions: (1) the effective temperature is *T*_*eff*_ ≈ *const*, (2) the volumetric strain energy increases with time as the consequence of normal residual stress accumulation, (3) the cell packing density increases with time, (4) internal entropy increases with time, i.e.*R*_*S*_ > 0, and (5) the effective chemical potential 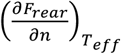 increases with cell packing density. Consequently, the volumetric strain energy density *W*_*V*_ accumulation induces an increase in the cell packing density per single viscoelastic state, expressed by eq. 14 as: 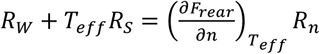. An increase in the cell packing density leads to a decrease in the cell mobility and on that base a decrease in the effective temperature. A decrease in the effective temperature perturbs the current equilibrium state, changes the state of viscoelasticity and on that base the rate *R*_*W*_. Five states of viscoelasticity gained within three regimes will be described by various constitutive models.

## 5. Viscoelasticity of multicellular systems for various regimes

The viscoelasticity of multicellular systems changes from a regime to regime by increasing the cell packing density accompanied with change of cell mobility. The convective regime accounts for two states of viscoelasticity: a viscoelastic liquid and viscoelastic solid, depending primarily on the magnitude of cell velocity and the state of cell-cell adhesion contacts (Pajic-Lijakovic and Milivojevic, 2021). Higher cell velocities and weak cell-cell adhesion contacts lead to pronounced liquid-like behavior while lower cell velocities and stronger cell-cell adhesion contacts ensure solid-like behavior (Pajic-Lijakovic and Milivojevic., 2019c;2021). Schematic presentation of viscoelastic states gained within the established regimes is shown in Figure 3.

**Figure 3.**
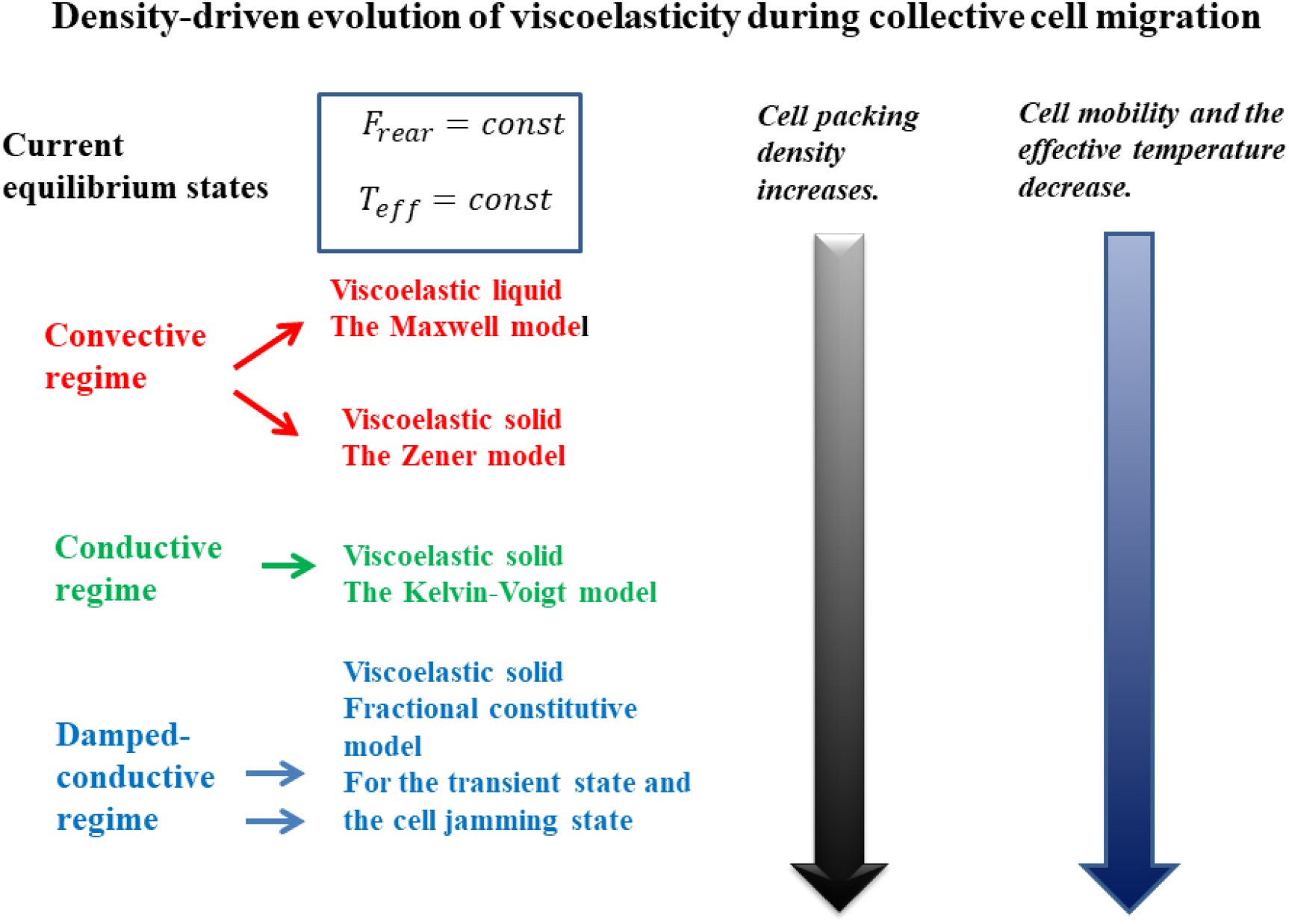
Various viscoelastic states gained within three regimes.

Guevorkian et al. (2011) considered micropipette aspiration of cell aggregate and proposed the Maxwell model suitable for a viscoelastic liquid. The average cell velocity is calculated as 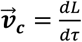 (where *L* is the length of the cell reached within a pipette). Corresponding average velocity in the initial regime is 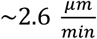 and decreases to 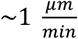 in the middle-time regime under the applied pressure of 500 *Pa*. Lee and Wolgemuth (2011) considered a long-time viscoelasticity caused by CCM of epithelial monolayer during wound closing. They also proposed the Maxwell model. Garcia et al. (2015) pointed out to the disordering trend of cell rearrangement which induces additional energy dissipation, characteristic for this state of viscoelasticity. This disordering was expressed by decreasing in the velocity correlation length with the cell velocity for 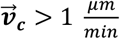.

Further increase in cell packing density within the same, convective regime leads to change the state of viscoelasticity from viscoelastic liquid to a viscoelastic solid. This viscoelastic solid state represents a characteristic of the confluent multicellular systems. Garcia et al. (2015) pointed out to the ordering trend of cell rearrangement characteristic for this state of viscoelasticity. This ordering was expressed by increasing in the velocity correlation length with the cell velocity for 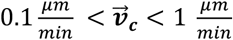. Notbohm et al. (2016) considered 2D cell swirling motion within a confluent monolayer by monitoring the maximum radial velocity of 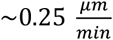. They reported that the long-time change of radial cell residual stress component 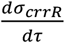 correlated well with the long-time strain change 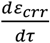 during the time period of 24 h. This result points out to the Zener constitutive model, suitable for viscoelastic solid, which was elaborated rheologicaly by Pajic-Lijakovic and Milivojevic (2020c).

The main characteristic of a viscoelastic liquid is that stress can relax under constant strain rate condition (the Maxwell model) (Pajic-Lijakovic, 2021). In the case of a viscoelastic solid, some linear models as the Zener model describes stress relaxation while the others as the Kelvin-Voigt model not. However, if the stress can relax, its relaxation occurs under constant strain condition (the Zener model). The stress relaxation has been monitored during cell aggregate uni-axial compression between parallel plates under constant aggregate shape (or surface strain) (Marmottant et al., 2009). The stress relaxation time is 3-14 min. Stress relaxed during 25 min from initial value to the residual value while the change of strain caused by CCM is much slower and corresponds to hours (Marmottant et al., 2009; Pajic-Lijakovic and Milivojevic 2020b;2021). During the established stress relaxation time of 10 min, cells are capable of moving along the distance equal to the single-cell size by the velocity of 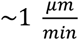 (the convective regime).

Transition from convective to the conductive regime is accompanied with significant reduction in cell mobility which causes an increase in the stress relaxation time. This so called “plateau” regime near the system jamming has been recognized for various soft matter systems and described by the Kelvin-Voigt model (Liu et al., 2017). The main characteristic of this regime is that stress cannot relax (Pajic-Lijakovic, 2021). Cells oscillate around their equilibrium states trapped into cages (Nnetu et al., 2013). Further increase in cell packing density leads to an anomalous nature of energy dissipation during cell long-time rearrangement characteristic for the damped-conductive regime (Nnetu et al., 2013). This regime accounts for the cell transient state and jamming viscoelastic states (Pajic-Lijakovic and Milivojevic, 2019c).

Consequently, five viscoelastic states gained within three viscoelastic regimes will be elaborated rheologicaly (Figure 3) by discussing the corresponding constitutive models such as: (1) the Maxwell model for a viscoelastic liquid (Guevorkian et al., 2011; Lee and Wolgemuth, 2011; Garcia et al., 2015) (for the convective regime: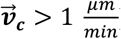), (2) the Zener model for a viscoelastic solid (Pajic-Lijakovic and Milivojevic, 2020c) (for the convective regime: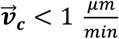), and (3) the Kelvin-Voigt model constitutive models for a viscoelastic solid (Liu et al., 2017) (for the conductive regime: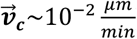), and (4) the fractional constitutive model for transient and jamming viscoelastic states (Pajic-Lijakovic and Milivojevic, 2019c) (for the damped-conductive regime: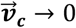).

### 4.1 The Maxwell model (the convective regime): for higher cell velocities

Single-cell movement within a stream induces generation of local shear strain rate 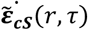 equal to 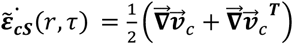 and local volumetric strain rate 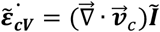 (where *Ĩ* is identity tensor) (Pajic-Lijakovic and Milivojevic, 2021). Both strain rates are supposed to be constant during a short-time relaxation cycle and changes from cycle to cycle. The short-time stress relaxation cycles correspond to the time scale of minutes (Marmottant et al., 2009; Pajic-Lijakovic and Milivojevic, 2019a;2021). These strain rates induce generation of corresponding shear and normal stresses and their relaxation during short-time relaxation cycles. We are interested in the accumulation of the normal residual stress and the volumetric strain energy density as a consequence of an increase in cell packing density (Trepat et al., 2009). The Maxwell model, in this case, is expressed as:

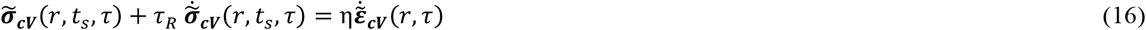

where 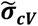 is the normal stress, 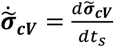, and η is the volumetric viscosity, *τ*_*R*_ is the corresponding stress relaxation time and, 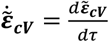 is the strain rate. Stress relaxation under constant strain rate 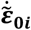 per single short-time relaxation cycle can be expressed starting from the initial condition 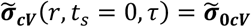 as:

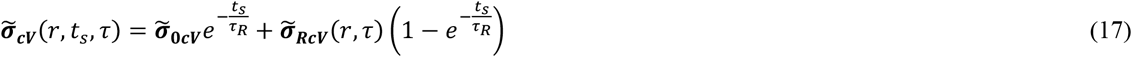

Where 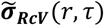 is the residual stress equal to 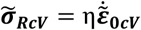. Residual stress accumulation can suppress cell migration and on that base change of the state of viscoelasticity. The corresponding rate of volumetric strain energy change (eq. 14) is 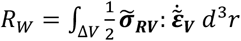.

Eq. 16 can be transformed from the time-domain into the frequency-domain using the Fourier integral transform. Transforming equation is expressed in the form *F*[σ (*t*)] = *G*^*^(*ω*)*F*[ε (*t*)] (where *F*[·] is the Fourier transform, *ω*is the angular velocity, and *G*^*^(*ω*) is the complex modulus) (Pajic-Lijakovic, 2021). The complex modulus is equal to: *G*^*^(*ω*) = *G*^’^(*ω*) + *i G*^″^(*ω*) (where *G*^’^(*ω*) is the storage modulus, *G*^″^(*ω*) is the loss modulus, and 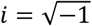 is the imaginary unit). The storage modulus *G*^’^(*ω*) quantifies elastic behaviour while the loss modulus *G*^″^(*ω*) quantifies viscous behaviour of the examined system. A higher value of the storage modulus indicates more intensive solid-like behaviour while a higher value of the loss modulus indicates more intensive liquid-like behaviour. The storage and loss moduli for the Maxwell model are (Pajic-Lijakovic, 2021):

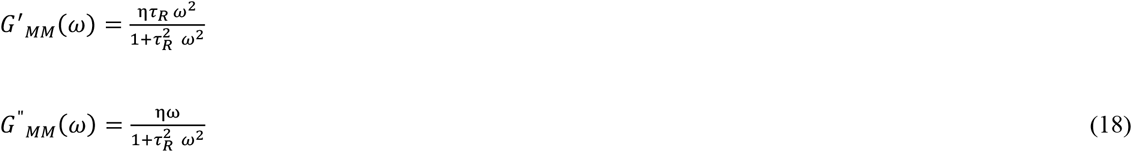

where *G*^’^_*MM*_(*ω*) is the storage modulus and *G*^″^_*MM*_(*ω*) is the loss modulus for the Maxwell model. Viscoelastic liquids described by the Maxwell model show: (1) liquid-like behaviour for lower angular velocities *G*^’^_*MM*_(*ω*)/ *G*^″^_*MM*_(*ω*) < 1 and (2) solid-like behaviour *G*^’^_*MM*_(*ω*)/ *G*^″^_*MM*_(*ω*) >1 for larger angular velocities (Pajic-Lijakovic, 2021). An increase in cell packing density reduces the cell mobility described by the effective temperature and perturbs the state of viscoelasticity (Figure 3). This perturbation leads to viscoelastic liquid-to viscoelastic solid cell state transition within the same convective regime. A new viscoelastic solid state corresponds to a new current equilibrium state and can be described by the Zener model (Pajic-Lijakovic and Milivojevic, 2020b,2021).

### 5.2 The Zener model (the convective regime): for lower cell velocities

Step by step movement of the strongly connected cell cluster through dense surrounding induces generation of local shear strain 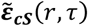 and volumetric strain 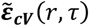 within the cell cluster. The volumetric compressive strain is also intensive during collision of velocity fronts caused by uncorrelated motility (Pajic-Lijakovic and Milivojevic, 2019a;2021). For small strain assumption these strains can be expressed 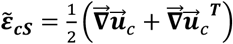 and 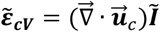. These strains are constant per single short-time relaxation cycle and change from cycle to cycle as a result of cell clusters movement (Pajic-Lijakovic and Milivojevic, 2019a;2020b). The strains induce generation of corresponding shear stress 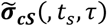 and normal (compressive/tensile) stress 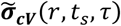. These stresses relax during short-time relaxation cycles up to residual values 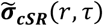 and 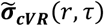, respectively. We are interested in the accumulation of the normal residual stress as a consequence of an increase in cell packing density. The simplest constitutive model of a viscoelastic solid capable for describing stress relaxation under constant strain condition is the Zener model (Pajic-Lijakovic and Milivojevic, 2020b;2021) expressed as:

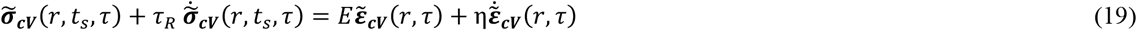

where 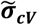 is the shear or volumetric stress, 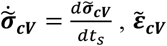 is the volumetric strain, 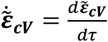 is the strain rate, *E* is the Young’s elastic modulus equal to 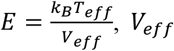 is the effective volume per single cell, and η is the volumetric viscosity equal to η = E *t*_*r*_,*t_r_* is the strain relaxation time under constant stress condition, and τ*_R_* is the stress relaxation time. The corresponding volumetric strain energy density is 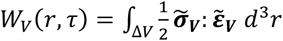.

Stress relaxation under constant strain condition 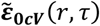 per single short-time relaxation cycle can be expressed starting from the initial condition 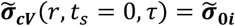 as:

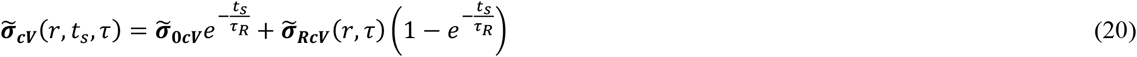

where 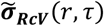 is the residual shear or normal stress equal to 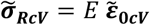. Notbohm et al. (2016) reported that the long-time change of radial cell residual stress component 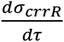 correlated well with the long-time strain change 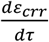 which represents the confirmation of the viscoelastic solid behavior described by the Zener constitutive model σ (Pajic-Lijakovic and Milivojevic, 2020c). Complex modulus *G*^*^(*ω*) can be formulated after Fourier transform of eq. 19. Corresponding storage and loss moduli can be simplified as (Pajic-Lijakovic, 2021):

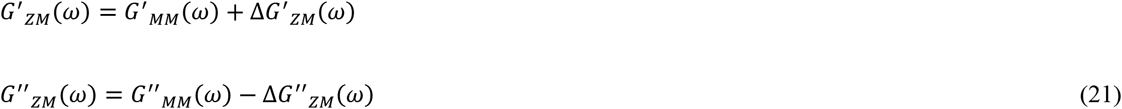

where *G*^’^_*ZM*_(*ω*) is the storage modulus and *G*^″^_*ZM*_(*ω*) is the loss modulus for the Zener model, *G*^’^_*MM*_(*ω*) is the storage modulus and *G*^″^_*MM*_(*ω*) is the loss modulus for the Maxwell model (eq. 18), Δ*G*^’^_*ZM*_(*ω*) is the contribution to storage modulus equal to 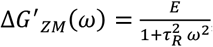, and *G*^″^_*ZM*_(*ω*) is the contribution to loss modulus equal to 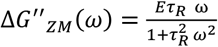. Storage modulus for the Zener model is larger than the one formulated for the Maxwell model (eq. 16). Loss modulus for the Zener model is lower than the one expressed for the Maxwell model. The deviation of *G*^’^_*ZM*_(*ω*) from *G*^’^_*MM*_(*ω*) is pronounced for lower angular velocities, while for higher angular velocities Δ*G*^’^_*MM*_(*ω*) → 0 (Pajic-Lijakovic, 2021).

The stress relaxation time increases with the cell packing density while the rate 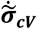 decreases as a consequence of the cell mobility decrease. When the rate 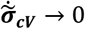, the transition from convective to the conductive regime occurs. For this condition the Zener model evolves into the Kelvin-Voigt model. Both models correspond to a viscoelastic solid state.

### 5.3 The Kelvin-Voigt model (the conductive-plateau regime) – for low cell velocities

The main characteristic of this viscoelastic state is the suppression of the stress relaxation, i.e 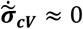. It represents a consequence of a decrease in the cell mobility (Figure 3). Corresponding rheological behavior can be described by the Kelvin-Voigt model. This model has been already applied for describing the viscoelasticity of various soft matter systems near jamming or glass transitions (Liu et al., 2017). The Kelvin-Voigt model describes stress change by changing the strain and strain rate as:

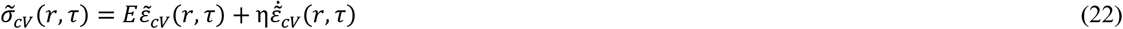

Where *E* is the Young’s elastic modulus equal to 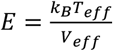, and is the volumetric viscosity equal to η = E *t*_*r*_, *t*_*r*_ is the strain relaxation time under constant stress condition, such that: (1) the effective volume *V*_*eff*_ ^*cd*^ < *V* _*eff*_ ^*cv*^, (2) the strain relaxation time *t*_*r*_ ^*cd*^ > *t*_*r*_ ^*cv*^, and (3) the effective temperature *T*_*eff*_ ^*cd*^ < *T* _*eff*_ ^*cv*^ .The normal stress is equal to 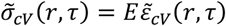 under constant strain condition. The corresponding volumetric strain energy is 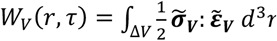. The storage and loss moduli, obtained after Fourier transform of eq. 22, are (Pajic-Lijakovic, 2021):

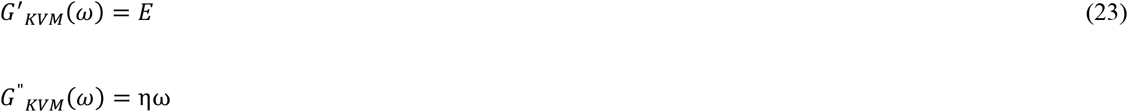

where *G*^’^_*KVM*_(*ω*) is the storage modulus and *G*^″^_*KVM*_(*ω*) is the loss modulus for the Kelvin-Voigt model. Constant value of the storage modulus *G*^’^(*ω*) obtained for the Kelvin-Voigt model represents the characteristic of this plateau regime. Viscoelastic solids described by Kelvin-Voigt model show: (1) solid-like behaviour for lower angular velocities *G*^’^_*KVM*_(*ω*)/ *G*^″^_*KVM*_(*ω*) > 1 and (2) liquid-like behaviour *G*^’^_*KVM*_(*ω*)/ *G*^″^_*KVM*_(*ω*)< 1 for larger angular velocities (Pajic-Lijakovic, 2021)

Further increase in cell packing density additionally reduced cell movement which leads to the anomalous nature of energy dissipation during cell rearrangement (Nnetu et al., 2013; Pajic-Lijakovic and Milivojevic, 2021). Transition from the conductive to the damped-conductive regime occurs for *T*_*eff*_ → 0 and consequently, for the Young’s modulus *E* → 0 under these conditions.

### 5.4 Constitutive model for the damped-conductive regime

The main characteristics of the damped-conductive regime is the anomalous nature of energy dissipation during cell rearrangement and the Young’s modulus *E* → 0. Consequently, the Kelvin-Voigt model proposed to the conductive regime evolves into a new, fractional constitutive model proposed by Pajic-Lijakovic and Milivojevic (2019c,2021):

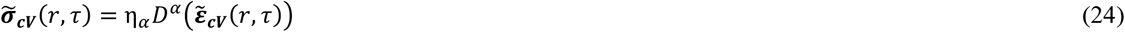

Where η_*α*_ is the effective modulus, while 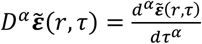 is the fractional derivative, and is the order of fractional derivative i.e. the damping coefficient of system structural changes which satisfies the conditions (1) *α* = 1/2 for the transient state and (2) *α* < = 1/2 for the jamming state (Pajic-Lijakovic and Milivojevic, 2019c,2021). Baumgarten and Tighe, (2017) pointed out that systems have to pass through the transient state before jamming. If the damping coefficient is *α* = 0, it is obtained that 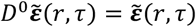 while the effective modulus tends to elastic modulus η_*α*_ → *E* (where is the Young’s modulus of elasticity). Higher values of the damping coefficient in the range 1> *α* > = 1/2 correspond to the viscoelastic liquid rather than viscoelastic solid (Pajic-Lijakovic, 2021). The corresponding rate of the volumetric strain energy change (eq. 14) is 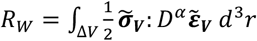. The storage and loss moduli for the damped-conductive regime can be expressed after Fourier transform of eq. 22 as (Pajic-Lijakovic and Milivojevic, 2019c):

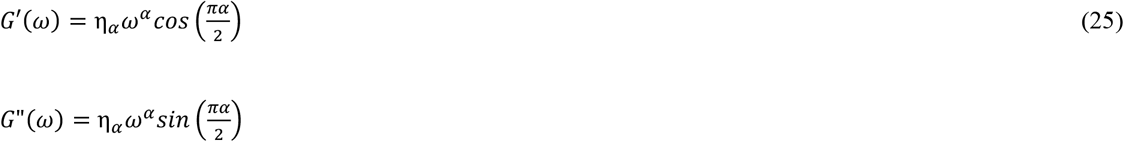

Where *α* = 1/2 and *G*^’^(*ω*) = *G*^″^(*ω*) corresponds to the transient state. Further reduction of cell mobility leads to cell jamming realized for *α* < = 1/2 and *G*^’^(*ω*) > *G*^″^(*ω*) which corresponds to the viscoelastic solid. Nnetu et al. (2013) reported that MCF-10A cell monolayers reaches jamming state for the cell packing density of 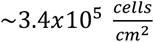. The corresponding damping coefficient obtained by monitoring mean square displacement vs. time is *α* ∼ 0.2 (Nnetu et al., 2013). Cells are capable of escaping the jamming state via CIL and start migrating again (Park et al., 2015). Nnetu et al. (2013) observed two jamming time-periods during 20 h of 2D CCM.

## 6. Conclusion

Significant attempts have been made to discuss the density-driven cell jamming state transition during 2D CCM. The cell jamming represents a consequence of the cell mobility reduction caused by the normal residual stress accumulation. The residual stress accumulation influences: (1) cell packing density, (2) cell−cell adhesion energy, (3) magnitude of cellular forces, (3) cell shape, (3) CIL and (4) their inter-relations. The reduction in cell mobility influences the state of viscoelasticity. Despite extensive research devoted to study of the jamming state transition we still do not understand the process from the standpoint of rheology.

This contribution represents an attempt to bridge this gap and clarify the relationship between viscoelasticity on one hand and cell packing density and cell mobility on the other. This density-driven evolution of viscoelasticity accounts for several transitions between five viscoelastic states gained within three regimes: (1) convective regime, (2) conductive regime, and (3) damped-conductive regime. The convective regime accounts for two states of viscoelasticity depending on the magnitude of cell velocity and the state of cell-cell adhesion contacts. Higher cell velocities and weak cell-cell adhesion contacts (characteristic for lower cell packing densities) lead to pronounced liquid-like behavior described by the Maxwell model, while lower cell velocities and stronger cell-cell adhesion contacts (characteristic for confluent multicellular systems) ensure solid-like behavior described by the Zener model. Transition from convective to the conductive regime is accompanied with significant reduction in cell mobility caused by increasing the normal residual stress accumulation. This so called “plateau” regime near the system jamming has been described by the Kelvin-Voigt model. Further increase in cell packing density leads to an anomalous nature of energy dissipation during CCM as a characteristic of the damped-conductive regime. This regime accounts for two states of viscoelasticity, i.e. the cell transient state and the jamming state described by the fractional constitutive model. The evolution of the viscoelasticity is realized through several current equilibrium states. Every equilibrium state was described thermodynamically by constant values of: (1) the Helmholtz free energy of cell rearrangement and (2) the effective temperature. The density-driven reduction of cell mobility perturbs the equilibrium state by decreasing the effective temperature. Next equilibrium state accompanied with a new state of viscoelasticity is established for the new values of the effective temperature and the Helmholtz free energy of rearrangement.

Additional experiments are necessary in order to (1) correlate the normal residual stress accumulation with change in the cell packing density, (2) identify the cell number density and the damping coefficient for the jamming state of various cell types, (3) find the corresponding parameters for the proposed constitutive models, (4) identify the time evolution of viscoelastic change from the convective regime to the damped-conductive regime under various 2D and 3D experimental conditions, and (5) find the necessary time period for cells to undergo unjamming transition.

## Acknowledgement

This work was supported by the Ministry of Education, Science and Technological Development of the Republic of Serbia (Contract No. 451-03-9/2021-14/200135)

## Notes

### Competing Interest Statement

The authors have declared no competing interest.

